# Diploid male gametes circumvent hybrid sterility between Asian and African rice species

**DOI:** 10.1101/2020.05.27.119180

**Authors:** Daichi Kuniyoshi, Itaru Masuda, Yoshitaka Kanaoka, Yuki Shimazaki-Kishi, Yoshihiro Okamoto, Hideshi Yasui, Toshio Yamamoto, Kiyotaka Nagaki, Yoichiro Hoshino, Yohei Koide, Itsuro Takamure, Yuji Kishima

## Abstract

In F_1_ hybrids of *Oryza sativa* (Asian rice) and *O. glaberrima* (African rice), heterozygosity leads to a complete gamete abortion because of allelic conflict at each of the 13 *hybrid sterility* (*HS*) loci. We systematically produced 19 plants from the F_1_ hybrids of both the rice species by the anther culture (AC) method. Five of the 19 interspecific hybrid plants were fertile and able to produce seeds. Unlike ordinal doubled haploid plants resulting from AC, these regenerated plants showed various ploidy levels (diploid to pentaploid) and different zygosities (completely homozygous, completely heterozygous, and a combination). These properties were attributable to meiotic anomalies in the interspecific hybrid F_1_ plants. Examination of the genetic structures of the regenerated plants suggested meiotic non-reduction took place in the interspecific hybrid F_1_ plants. The centromeric regions in the regenerated plants revealed that the abnormal first and/or second divisions of meiosis, namely the first division restitution (FDR) and/or second division restitution (SDR), had occurred in the interspecific hybrid. Immunohistochemical observations also verified these phenomena. FDR and SDR occurrences at meiosis might strongly lead to the formation of diploid microspores. The results demonstrated that meiotic anomalies functioned as a reproductive barrier occurred before the *HS* genes acted in gamete of the interspecific hybrid. Although such meiotic anomalies are detrimental to pollen development, the early rescue of microspores carrying the diploid gamete resulted in the fertile regenerated plants. The five fertile plants carrying tetraploid genomes with heterozygous alleles of the *HS* loci produced fertile diploid pollens, implying that the diploid gametes circumvented the allelic conflicts at the *HS* loci. We also proposed how diploid male gametes avoid HS with the killer-protector model.

## Introduction

Although cultivated rice species *Oryza sativa* (Asian rice) and *O. glaberrima* (African rice) both have AA genomes, the first filial generation (F_1_) between these two species does not produce fertile seeds [1, 2]. This type of reproductive isolation, designated as hybrid sterility (HS), is associated with abnormal gamete development and sterility [1, 2, 29]. To date, 13 *HS* loci have been reported to be involved in HS in F_1_ hybrids between *O. sativa* and *O. glaberrima* (*sat–gla*) [4-7, 9-15]. In particular, pollen sterility is noticeable in these hybrids, and fertility is completely lost; in contrast, female gametes do not exhibit such severe sterility, as seeds are produced when fertile pollen grains are crossed [16, 17]. However, microspores in the process of completing meiosis and developing into pollen can differentiate into plants. If pollen destined for abortion can be rescued during early developmental stages, it could create hybrid plants. Not only are these individuals useful as genetic resources, but they also have a high potential in elucidating the mechanism of hybrid sterility.

In the 1960s, Gopalakrishnan et al. [30] and Oka [31] created individuals producing fertile seeds in F_1_ tetraploid hybrids of *sat–gla*. In 1980, Woo and Huang reported that anther culture (AC) of an F_1_ hybrid of *sat–gla* gave rise to tetraploid, diploid, and haploid plants [32]. Unfortunately, these significant findings were given scant attention, being published too early to be of wide interest. The results described in those studies have thus not been validated, and the fertility of such F_1_ tetraploid hybrids has not been analyzed in detail. Furthermore, the mechanism responsible for the formation of polyploids following AC of the interspecific hybrids has not been studied subsequently. In these interspecific hybrids, detailed observations are required to determine if pairing between genomes occurs during meiosis and whether distributions of homologous chromosomes in the first meiotic division and/or sister chromatids in the second meiotic division take place.

AC technology developed in the 1960s [20, 21] and now widely used in crops [22, 23], haploid genomes derived from male gametes can be doubled to form a doubled haploid (DH) individual with complete homozygosity. The differentiated individual from AC is, therefore, a complete pure line. In autogamous crops, a pure line is an essential condition for cultivars; consequently, AC technology has contributed to the breeding of many crops [24, 25].

Our previous study revealed that AC of an F_1_ hybrid of *sat–gla* can generate calli from anther-containing microspores in the late uninucleate stage [26]. In the present study, 19 plants were differentiated from such calli after a regeneration treatment, and some individuals of successively obtained plants produced seeds. The differentiated individuals were tetraploid and exhibited heterozygosity in many genomic regions, which might cause allelic conflicts of *HS* loci. Many studies on HS have mainly focused on *HS* genes such as HS loci, but little attention has been paid to other genetic factors. These tetraploids were mainly a consequence of meiotic anomalies attributable to a failure during first or second meiotic divisions. Here, we demonstrate that diploid gametes can circumvent HS between *sat–gla* and thus allow fertile individuals to be regenerated. We also examine the relationship between meiotic anomalies and HS and discuss the defeat of HS by polyploidization.

## Results

### Pollen sterility of interspecific F_1_ hybrids

Interspecific hybrids between *sat–gla* are well known to exhibit severe HS that possibly involves more than a dozen *HS* genes (Sano et al., 1979; Sano, 1983, 1990; Doi et al., 1998; Doi et al., 1999; Taguchi et al., 1999; Ren et al., 2006; Zhang et al., 2006; Li et al., 2011; Xu et al., 2014; Yu et al., 2018). An interspecific F_1_ hybrid of *O. sativa* L. ssp. *japonica* Nipponbare (Nip) and *O. glaberrima* Steud. accession IRGC 104038 from Senegal (designated as WK21) produced panicles with sterile seeds as a consequence of aborted pollen and a partially fertile embryo sac (Fig. 1A). The mature pollen grains from WK21/Nip F_1_ were less strongly stained by Lugol’s solution, indicating their sterility and inability to accumulate polysaccharides (Fig. 1A). To explore the progression of this pollen sterility, we observed developing pollen grains in Nip, WK21, and WK21/Nip F_1_ (Fig. 1B). As development continued, pollen grains of both parents first showed evidence of acetocarmine staining at the early binucleate stage and were fully stained at the trinucleate stage (Fig. 1B). During the early uninucleate stage of pollen development, most microspores from WK21/Nip F_1_ plants exhibited no prominent differences in size or shape compared with the parents (Fig. 1B), but some had abnormal structures, such as a fused form or a larger size than that of normal microspores (Fig. 1C). The proportion of standard-shaped microspores in WK21/Nip F_1_ plants decreased as they developed into pollen (Fig. 1B). At the mature stage, normal, round pollen grains had disappeared, and the number of cavitated pollen grains had increased (Fig. 1B).

**Fig. 1.**
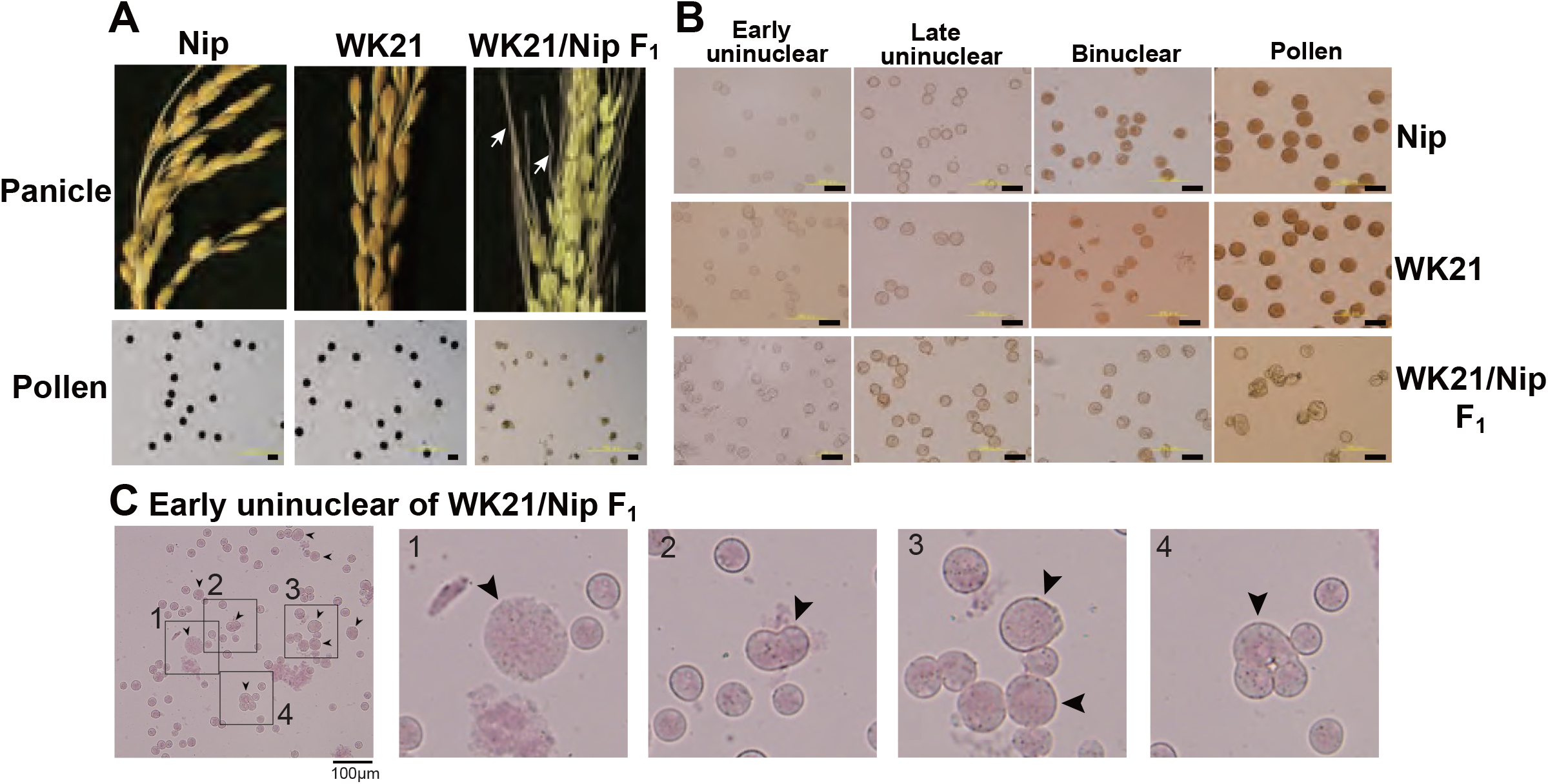
Images of microspores at different developmental stages in Nip, WK21, and WK21/Nip F_1_ plants. (A) Panicles and pollen grains of Nip, WK21, and WK21/Nip F_1_ plants. Mature panicles were observed in individuals of Nip, WK21, and their interspecific F_1_ hybrid (WK21/Nip F_1_) at the ripening stage. Panicles in Nip and WK21 were fertile, while the panicle in WK21/Nip F_1_ was sterile. Awns developed in the interspecific F_1_ hybrid (white arrows) but not in the parents. All plant materials used for experiments were grown in a greenhouse. Pollen grains from Nip and WK21, which were stainable with Lugol’s iodine solution, exhibited potential fertility, whereas pollen from WK21/Nip F_1_ was sterile, as reflected by the absence of staining. (B) Microspores at early uninucleate, late uninucleate, binucleate, and trinucleate stages. Microspores were stained with acetocarmine. The black bar in each panel corresponds to 100 μm. WK21/Nip F_1_ plants at the early uninucleate stage seemed to contain mostly normal microspores. The number of abnormal microspores increased as development progressed until most pollen grains appeared cavitated. (C) Abnormal microspores in WK21/Nip F_1_ plants at the early uninucleate stage. Microspores were stained with acetocarmine. Black arrows indicate abnormally shaped microspores. The black bar in each panel corresponds to 100 μm.

### Plant regeneration from calli of interspecific F_1_ hybrids

In previous research, Kanaoka et al. (2018) successfully rescued microspores at the late uninucleate stage in interspecific hybrid plants (WK21/Nip F_1_ and its reciprocal cross hybrid Nip/WK21 F_1_) to induce calli by the AC method with RI-13 medium. In that study, 98 calli were obtained from 28,181 anthers, which corresponded to induction frequencies of approximately 11 calli from 14,724 Nip/WK21 anthers and 87 calli from 13,457 WK21/Nip anthers (Supplemental Table S1). In the present study, we used the 87 calli derived from WK21/Nip F_1_ for plant regeneration. The 11 Nip/WK21 calli (the opposite cross combination to WK21/Nip) were not used because only a single plantlet was generated. Distinct frequencies of callus generation between the two reciprocal hybrids were used to infer whether certain sporophytic influences were due to cytoplasmic or maternal effects of the parental plants. Regeneration of plants from calli was attempted using N6-based medium. We obtained 19 regenerated plantlets from the WK21/Nip F_1_-anther derived calli (Supplemental Table S1; Supplemental Fig. S1). Thirteen plantlets were regenerated from 23 Nip calli induced with SK-1 medium, whereas no plantlets were regenerated from WK21 calli in this study (Supplemental Table S1). The 19 plantlets from the WK21/Nip F_1_-anther derived calli were grown in soil; 17 became mature plants, while two died (Supplemental Fig. S1). Two phenotypic traits typically different between *sat–gla*, namely, leaf smoothness and awn presence, were segregated in the 17 regenerated plants and both parents (Fig. 2A, B).

**Fig. 2.**
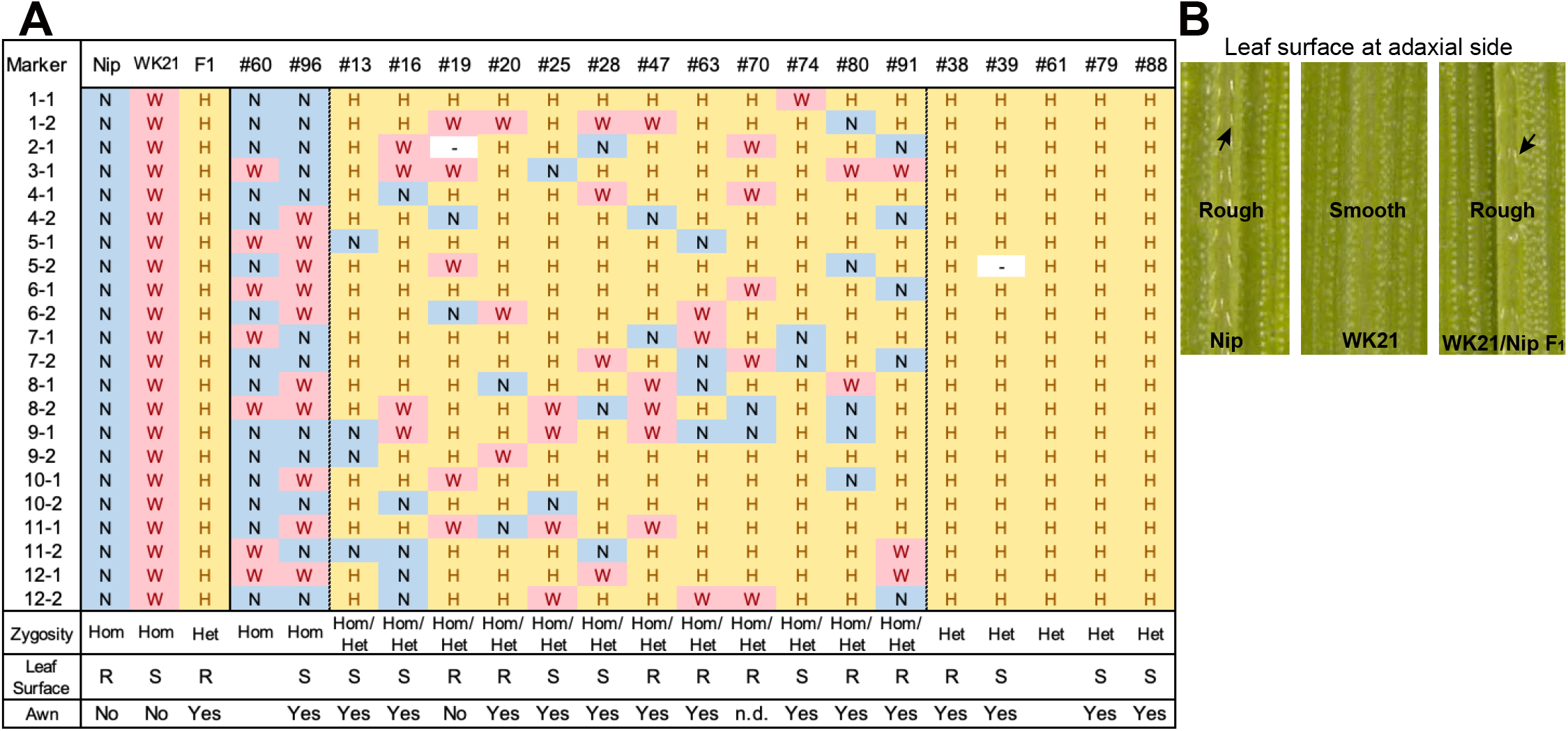
Characteristics of the 19 regenerated plants. (A)Genotypes of 19 plants regenerated from the calli of WK21/Nip F_1_ hybrids. The markers used for genotyping—one or two selected from each of 12 chromosomes—are detailed in Supplemental Fig. S2. Two regenerated plants, #60 and #96, were homozygous in all marker regions (Hom). The next 12 regenerated plants contained both homozygous and heterozygous regions (Hom/Het). The five plants on the far right-hand side were determined to be heterozygous (Het). At each marker position, the presence of two *Oryza sativa* (Nip) alleles, two *O. glaberrima* (WK21) alleles, or one copy of each (heterozygous) is indicated in the table by “N”, “W”, and “H”, respectively. The state of the leaf surface [rough (R) or smooth (S)] and the presence (yes) or absence (no) of awns (Fig. 1A) are also indicated. (B) Leaf surfaces of Nip, WK21, and WK21/Nip F_1_ plants. Surfaces of adaxial sides of Nip and F_1_ leaves were rough because trichomes were present (black arrows), whereas those of WK21 were smooth because trichomes were lacking.

### Genotyping of regenerated plants

The 19 plantlets grown as seedlings from callus were genotyped with 22 simple sequence repeat (SSR) markers located on the 12 rice chromosomes and polymorphic between the two parents (Supplemental Fig. S2, S3). In general, the DH plants obtained via AC had completely homozygous genomes as a result of the doubling of the male gametic genome. Any heterozygotes may have been due to DNA of somatic tissues (e.g., from anther walls) of the F_1_ hybrid plants, but we could not rule out the possibility of allopolyploids involving both parental genome sets. As shown in Fig. 2A, genotyping of the 19 plantlets revealed that two plantlets (#60 and #96) were completely homozygous (Hom) for either genotype at each marker locus, while five plantlets (#38, #39, #61, #79, and #88) were heterozygous (Het) at all loci. The remaining 12 individuals had mixed genomes (Hom/Het) containing both homozygous and heterozygous loci (Fig. 2A). The coexistence of homozygous and heterozygous loci in the plantlets derived from AC has two possible causes: an abnormality of meiosis in the parental plants or fusions between cells containing homozygotes and/or heterozygotes during callus culture. These results are in contrast to the observations of Morinaga and Kuriyama (1957), who did not detect any meiotic anomalies in their cytological study of interspecific hybrids between *sat–gla*.

### Ploidy analysis of regenerated plants

To examine ploidy levels of the 19 regenerated plantlets obtained from AC, we performed a flow cytometric analysis (Fig. 3A). Ploidy levels of the analyzed samples were based on relative fluorescence intensity comparisons with the parental diploid. As shown in Fig. 3B, five of the 19 plantlets were diploid, and 12 regenerated plants—eight tetraploids, three triploids, and a pentaploid—were polyploid. No haploids were obtained. No apparent relationship was observed between ploidy level and degree of homo- or heterozygosity, but the three triploids were commonly Het plantlets (Fig. 2; Supplemental Table S2). Among the 12 Hom/Het plantlets, five were diploid, one was pentaploid, and six were tetraploid (Supplemental Table S2). Microscopic observation also supported the results of the flow cytometric analysis: root tip cells from the examined plantlet (#20) had a chromosome number larger than 40, compared with 24 chromosomes in the parental *sat–gla* diploid (Fig. 3C). Unlike AC of intraspecific hybrids, which usually produces DH plants, AC of the interspecific *sat–gla* F_1_ hybrid resulted in many polyploid regenerants (12/19). These results led us to consider whether microspores from the F_1_ hybrid were directly responsible for the aberrant ploidy levels.

**Fig. 3.**
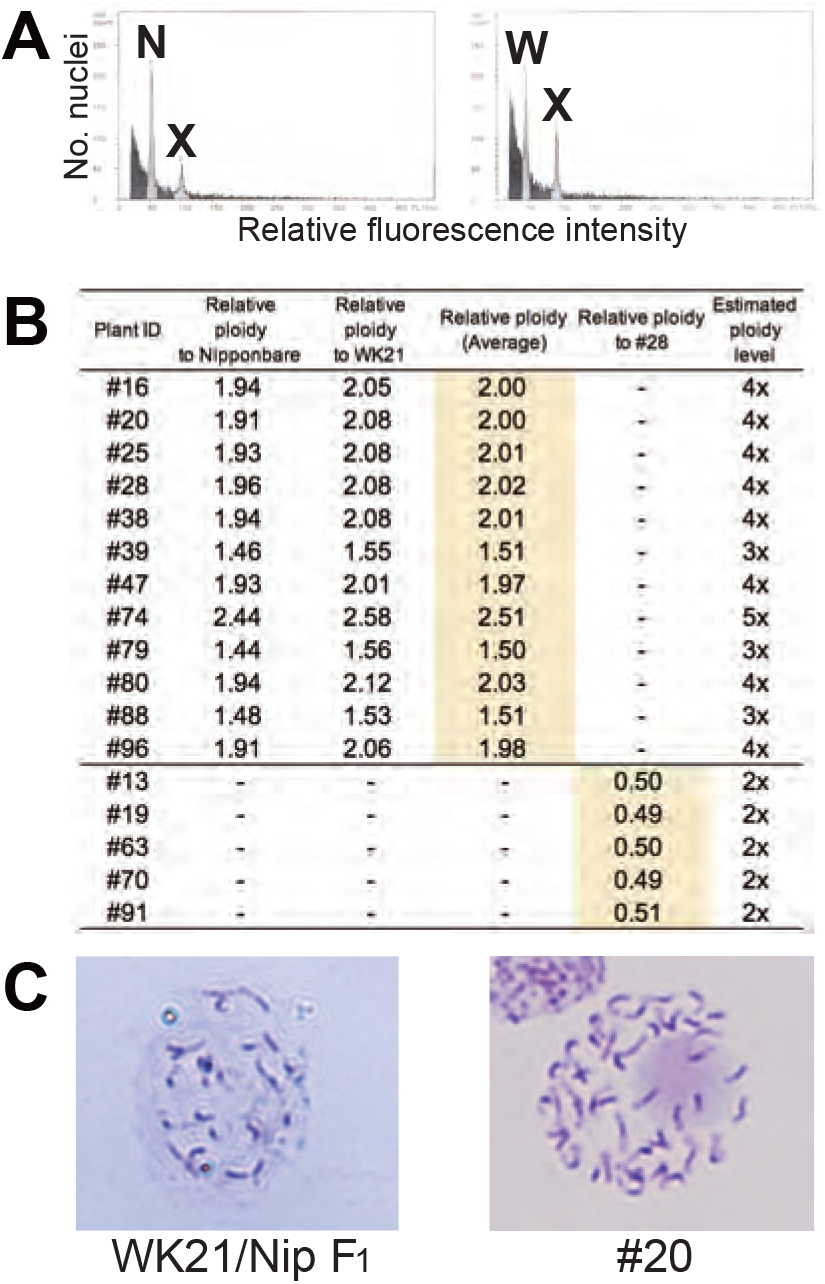
Ploidy analyses of somatic cells of regenerated plants based on flow cytometry (FCM) and Giemsa staining. (A) FCM-based ploidy analysis. Left: FCM histogram of samples of Nip and plant #20 showing two peaks—N (derived from the Nip genome) and X (derived from the #20 genome). Right: FCM histogram showing two peaks—W (from WK21) and X. The ploidy level of #20 was determined by comparing peak X with peaks N and W from the diploid parental lines. The relative fluorescence intensity of the peak of #20 was nearly twice as high as that of the two parents. (B) Ploidy levels of regenerated plants estimated from fluorescence intensity peak ratios. Ploidy levels of regenerated plants were based on relative fluorescence intensities of nuclei in Nip and WK21 cells. Diploid ploidy levels estimated by this method were validated by comparison with regenerated plant #28, which was determined to be tetraploid. (C) Giemsa staining of mitotic cells. Left: mitotic cell from the root tip of a WK21/Nip F_1_ plant. The number of chromosomes in the cell appears to be a half that of a #20 plant. Right: mitotic cell from the root tip of a #20 plant regenerated from WK21/Nip F_1_. More than 40 chromosomes are visible.

### Origin of the Hom/Het plants

We considered three possible causes for the polyploidy of the regenerants. First, the 12 Hom/Het plants (#13, #16, #19, #20, #25, #28, #47, #63, #70, #74, #80, and #91) were expected to result from the generation of abnormal tetrads through incomplete meiotic reduction. These meiotic anomalies involve two major arrests of meiotic reduction: first division restitution (FDR) and second division restitution (SDR) (Jauhar, 2007; De Storme and Geelen, 2013; Han et al., 2018) (Fig. 4A). FDR is the halt in division of homologous chromosomes after recombination during meiosis I, while SDR is the arrest of the separation of paired sister chromatids during meiosis II (Fig. 4A). Either meiotic division restitution produces microspores carrying diploid Hom/Het genomes. Diploid microspores with Hom/Het genomes may be duplicated during callus formation or regeneration processes, resulting in tetraploid Hom/Het plants. In regard to possible causes of incomplete meiotic reduction, we could test whether FDR or SDR was responsible for the Hom/Het plants. Hom/Het plants arising by FDR were expected to exhibit heterozygosity (i.e., both parental sequences) around centromeric regions (De Storme and Geelen, 2013). Because centromeric regions rarely undergo recombination, centromeric regions in paired homologous chromosomes between *sat–gla* remained heterozygous after meiosis I (Fig. 4A). In contrast, Hom/Het plants generated by SDR would have homozygous centromeric regions (i.e., either parental sequence) because of the cancellation of sister-chromatid separation during meiosis II (De Storme and Geelen, 2013) (Fig. 4A). To distinguish between these two possibilities, the 12 chromosomes of the 12 regenerants were genotyped using centromeric-region-specific SSR and insertion/deletion polymorphism (InDel) primers (McCouch et al., 2002) (Supplemental Fig. 2). Genotyping of the centromeric regions yielded homozygous bands for the Hom plants and heterozygous bands for the Het plants (Supplemental Table S2 and Fig. 3). Genotyping of the 12 Hom/Het plants uncovered two clear patterns: eight individuals (#13, #19, #20, #25, #47, #63, #74, and #80) were heterozygous for all the markers in centromeric regions, while the remaining four individuals (#16, #28, #70, and #91) were homozygous (Fig. 4B). These results suggest that the first eight Hom/Het plants resulted from FDR and that the latter four plants were derived from SDR. These observations of pollen mother cells (PMCs) verify the occurrence of abnormalities at meiosis in the interspecific F1 hybrid between *sat–gla* (Fig. 4C, D) possibly associated with the unusual shapes of microspores shown in Fig. 1C. Normal bivalent chromosomes observed at diplotene in meiosis I are necessary for reduction division, which leads to meiosis II, whereas univalent chromosomes in meiosis I are unable to undergo normal division, resulting in loss of meiosis I. We obtained evidence that germinal cells in the interspecific F_1_ hybrid retained univalent chromosomes. As shown on the left side of Fig. 4C, immunochemical staining with anti-*Oryza sativa* centromeric histone H3 (OsCenH3) antibody revealed a numerous pairs of centromeric signals, which implies alignment of bivalent chromosomes at diplotene in PMCs. In the same PMC sample, another gamete cell exhibited unpaired centromeric signals that were given by the presence of univalent chromosomes (right side of Fig. 4C). During anaphase II, we also observed unequal division, in which spindle fibers with α-tubulin were not equally formed in dividing cells (right side of Fig. 4D) relative to normal division (left side of Fig. 4D). These observations in PMCs of the interspecific F_1_ hybrid support the occurrence of FDR and SDR in meiosis I and II, respectively.

**Fig. 4.**
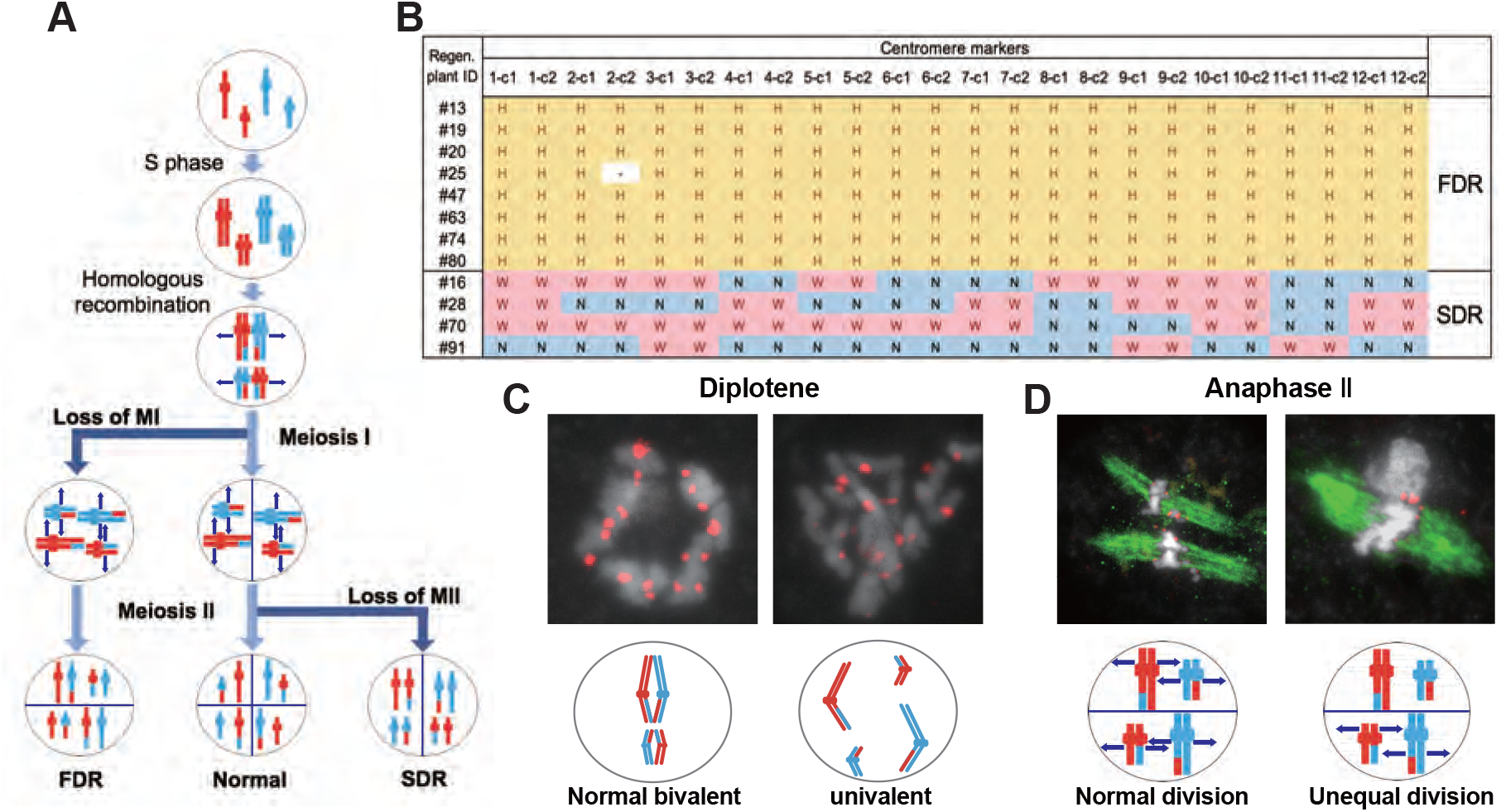
Meiotic anomalies associated with FDR and SDR in WK21/Nip F_1_. (A) Schematic diagram of chromosomal separations following normal, FDR, and SDR meiotic events. The three different chromosomal separation pathways following normal, FDR, and SDR events during meiotic division are based on De Storme and Geelen (2013). In the normal situation, bivalent homologous chromosomes separate after recombination at the end of meiosis I, with the sister chromatids remaining attached at the beginning of meiosis II and then separating. In FDR, homologous chromosomes fail to separate at the end of meiosis I, resulting in homologous chromosomes in the gametes. SDR bypasses meiosis II, and sister chromatids are distributed into gametes. FDR and SDR lead to centromeric regions (shown as knobs) that are respectively heterozygous or homozygous between homologous chromosomes. Red and blue are used to indicate the parental origin of chromosomal regions. (B) Genetic zygosities of centromeric regions of the 12 chromosomes of 12 regenerated plants and detection of FDR and SDR. Markers in centromeric regions used in this analysis are detailed in Supplemental Fig. S2. The 12 regenerated Hom/Het plants retaining both homozygous and heterozygous regions constitute two groups: those with completely heterozygous centromeric regions (#13 to #80) and those with completely homozygous ones (#16 to #91). The former group reflects the genetic nature of FDR, while the latter group is indicative of SDR. In the table, the presence at a given marker position of two *O. sativa* (Nip) alleles, two *O. glaberrima* (WK21) alleles, or one allele of each (heterozygous) is indicated by “N”, “W”, and “H”, respectively. (C) Immunohistochemical detection of normal and anomalous gametes during meiosis I in WK21/Nip F_1_. Using anti-OsCenH3 antibody, centromeric regions were observed in chromosomes at diplotene in meiosis I in PMCs from WK21/Nip F_1_. Left: detection of paired signals (red spots) from centromeres at diplotene in a PMC, implying normal bivalent chromosomes (white portions). Right: non-aligned, dispersed centromeric signals, indicative of univalent chromosomes. (D) Immunohistochemical detection of normal and anomalous gametes during meiosis II of WK21/Nip F_1_. Using anti-α-tubulin mouse antibody, spindle fiber formation (green zone) was observed at anaphase II in PMCs from WK21/Nip F_1_. During normal anaphase II, sister chromatids (white zone) prepared to move toward opposite poles of the cell to generate haploid gametes. Left: normal division, showing movement of sister chromatids to the poles via the spindle fibers in both compartments as monitored using α-tubulin antibody. Right: unequal division in a PMC. In the upper compartment, no α-tubulin was observed, and sister chromatids were unable to separate and move to the poles; in contrast, the movement of sister chromatids along spindle fibers was apparent in the lower compartment.

Second, five Het plants corresponding to three triploids (#39, #79, and #88), one tetraploid (#38), and one missing (#61) obviously contained both parental genomes (Fig. 2A). PMCs that failed to undergo both divisions at meiosis I and II may not have formed tetrads. The occurrence of both division restitutions in a single meiocyte may therefore have given rise to tetraploid Het plants; however, making an assumption about whether the heterozygotic status of the triploids was due to simple aberrant meiosis or a complex process mediated by other factors is difficult. Third, in the Hom plants (#60 and #96), #96 with tetraploid genome may have arisen by haploid gamete doubling, but we could not ascertain exactly when doubling occurred during the AC procedure (Fig. 2A).

### Fertility and *HS* locus genotypes of regenerated plants

Among the 19 plantlets obtained from AC, 17 grew to maturity, while two (#60 and #61) died at the seedling stage. Of the surviving regenerated plants, the five tetraploid ones (#20, #25, #38, #47, and #80) generated seeds (Table 1). More specifically, the five fertile tetraploid plants comprised four Hom/Het plants and one Het plant (Fig. 2A). The precise fertility of each of these five plants could not be determined because they had inherited the shattering trait from their parent *O. glaberrima* WK21; however, two regenerated plants, #38 and #80, produced a relatively higher number of seeds. To confirm that HS had been overcome in the regenerants, the 17 regenerants were genotyped using 12 SSR primers linked to known *HS* loci (Kanaoka et al., 2018). As shown in Table 1, plant #96 was homozygous for alleles from either of the two parents at each SSR locus (Table 1). In the four Het plants (#38, #39, #79, and #88), the *HS* locus-specific SSR markers were all heterozygous (Table 1). The Hom/Het plants were mixed, carrying both homozygotic and heterozygotic loci (Table 1). The three fertile tetraploid plants, #25, #38, and #80, were heterozygous at more than eight *HS* loci, a situation that would have caused sterility if these plants had been diploid. Our results are in agreement with the observations of Gopalakrishnan et al.(1964) and Oka (Oka, 1968) that the tetraploidy of the interspecific hybrid allowed escape from HS.

**Table 1.** Genotyping of the 12 *HS* loci in the 17 regenerated plants

| *HS* locus | #13 | #19 | #63 | #70 | #91 | #39 | #79 | #88 | #16 | #20 | #25 | #28 | #38 | #47 | #74 | #80 | #96 |
| --- | --- | --- | --- | --- | --- | --- | --- | --- | --- | --- | --- | --- | --- | --- | --- | --- | --- |
| *S*<sub>1</sub> | H | H | H | W | H | H | H | H | H | H | H | H | H | H | H | H | W |
| *S*<sub>3</sub> | H | H | H | N | W | H | H | H | N | H | H | N | H | W | H | H | W |
| *S*<sub>18</sub> | H | H | H | W | N | H | H | H | H | H | H | W | H | H | H | N | W |
| *S*<sub>19</sub> | H | H | H | N | W | H | H | H | H | H | H | H | H | H | N | W | W |
| *S*<sub>20</sub> | H | H | W | H | N | H | H | H | H | H | H | H | H | H | N | H | N |
| *S*<sub>21</sub> | H | H | N | W | N | H | H | H | H | N | W | H | H | H | N | H | N |
| *S*<sub>29(t)</sub> | H | H | H | H | H | H | H | H | H | H | H | H | H | H | N | N | W |
| *S*<sub>34(t)</sub> | H | W | N | H | W | H | H | H | W | H | N | H | H | H | H | H | N |
| *S*<sub>36(t)</sub> | H | H | H | H | H | H | H | H | H | H | H | H | H | H | H | H | W |
| *S*<sub>37(t)</sub> | H | H | H | W | N | H | H | H | W | H | H | W | H | H | H | H | N |
| *S*<sub>38(t)</sub> | H | H | H | W | H | H | H | H | N | N | H | W | H | H | H | H | N |
| *S*<sub>39(t)</sub> | H | N | H | H | N | H | H | H | H | N | N | H | H | N | H | N | N |
| Collected seeds no. | 0 | 0 | 0 | 0 | 0 | 0 | 0 | 0 | 0 | 2 | 4 | 0 | 18 | 1 | 0 | 32 | 0 |
| Ploidy | 2x | 2x | 2x | 2x | 2x | 3x | 3x | 3x | 4x | 4x | 4x | 4x | 4x | 4x | 5x | 4x | 4x |

**Table 2.**
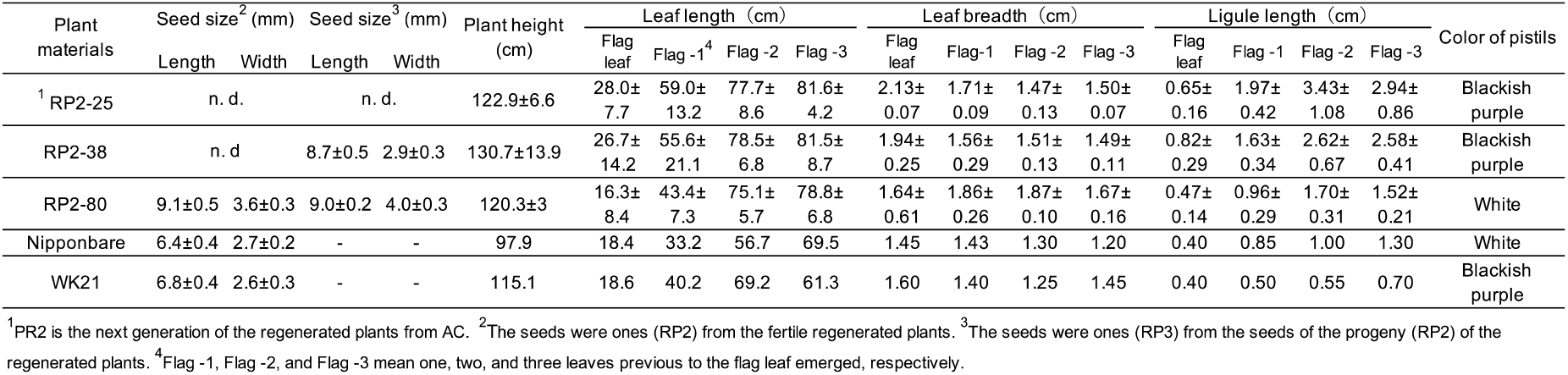
The features of the progenies (RP2-25, −38, −80) from the fertile regenerated plants, 25, 38, and 80, respectively

### Phenotypes of fertile tetraploids

Seeds (RP2) from #80 plants were larger than those of the parents: 1.42- and 1.34-fold longer and 1.33- and 1.38-fold wider relative to Nip and WK21, respectively (Supplemental Table S3). Likewise, fertile plants in the next generation (RP3) also produced bigger seeds (Supplemental Table S3). Of the five fertile tetraploid plants, the fertilities of plants from #25, #38, and #80 were passed along to subsequent generations. These plants thus appeared to have overcome the HS between *sat–gla*. Self-pollinated progenies of three fertile tetraploid lines, RP2-25 (from #25), RP2-38 (from #38), and RP2-80 (from #80), were obtained and phenotypically compared with their parental lines, Nip and WK21. The following characters were measured: seed length, seed width, plant height, leaf length, leaf breadth, ligule length, and pistil color (Supplemental Table S3). Most phenotypes in the second generation derived from the three tetraploid regenerated lines were larger than those of their parents (Supplemental Table S3), thus reflecting typical tetraploid vigor. In regard to pistil color, the blackish purple pistils of WK21 were expressed in the F_1_ generation and RP2-25 and RP2-38 lines, while the white pistils of Nip were inherited by the RP2-80 line (Supplemental Table S3).

## Discussion

### Production of plants from hybrids between *sat–gla* by AC

Because of gamete sterility, progenies cannot be generated from interspecific hybrids of *sat–gla* (Oka, 1957; Sano et al., 1979). In this study, we successfully regenerated plants from callus induced by culturing sterile microspores of interspecific hybrid plants without the recombinant DNA techniques. Five of the 19 regenerated plants produced seeds. According to Kanaoka et al. (2018), the essential factor for obtaining plants from interspecific hybrids with strong HS is the use of callus obtained by culturing anthers with microspores at the uninucleate stage. Uninucleate-stage microspores are required for embryogenesis not only in rice but also in wheat (Hassawi and Liang, 1990). In grape (Gribaudo et al., 2004), barley (Hoekstra et al., 1992), and *Brassica napus* (Telmer et al., 1992), embryoid bodies can also be differentiated directly from uninucleate microspores. Microspores appear to lose their embryogenic (or callus formation) ability after the uninucleate stage, and differentiation into pollen then irreversibly progresses (Kinoshita et al., 2000). Even in the interspecific hybrid between *sat–gla* exhibiting HS, microspore decay had not yet begun in uninucleate microspores (Figs. 1B, C, 5A). This stage is a crucial point for rescuing microspores to obtain plants from AC of interspecific hybrids (Kinoshita et al., 2000) (Fig. 5A).

**Fig. 5.**
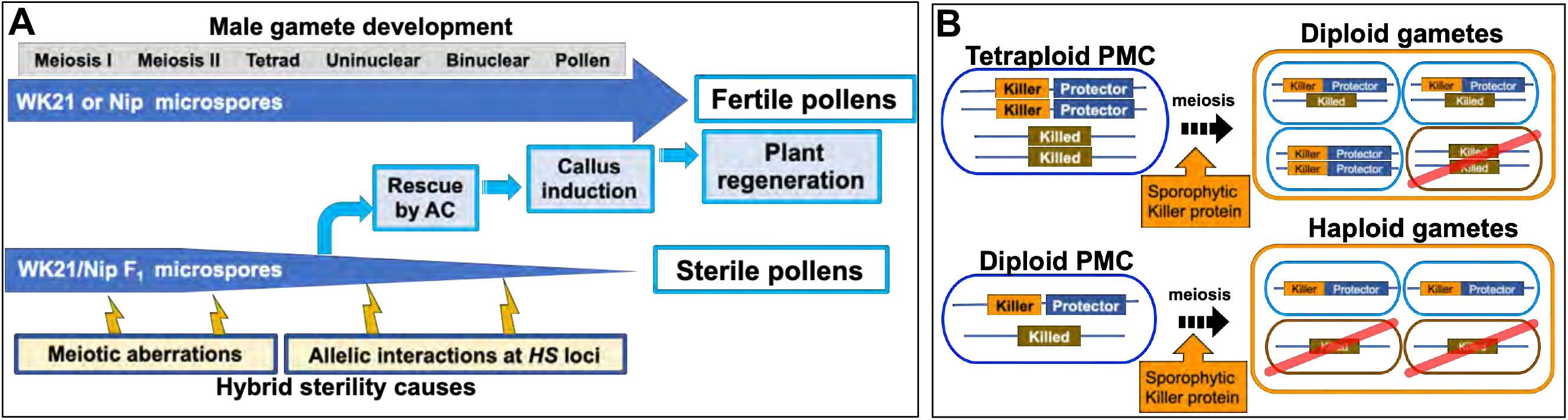
Models of processes of hybrid sterility and its circumvention. (A) A new model of hybrid sterility and plant regeneration by AC during microspore development in WK21/Nip F_1_. Meiotic aberrations are proposed as a cause of hybrid sterility. The parental varieties, WK21 and Nip, undergo normal microspore development to form pollen. In contrast, microspores of WK21/Nip F_1_ do not develop into pollen because of HS due to 1) meiotic aberrations and 2) allelic interactions at *HS* loci. (B) Higher rates of fertile gametes in tetraploids compared with diploids according to the killer–protector model. Under the killer–protector model of HS, the killer protein has a sporophytic effect on gametes during microspore development after meiosis, but the gamete expressing the protector protein is not killed. In the case of a heterozygous tetraploid plant, which contains two killer–protector alleles and two killed alleles, three-quarters of the gametes possess protector alleles. In the case of a heterozygous diploid plant, only half of the gametes carry a killer–protector allele. In theory, a heterozygous tetraploid thus produces 25% more surviving gametes than does a heterozygous diploid plant.

Diploid plants differentiated through AC usually have complete homozygosity because haploid male gametes are spontaneously doubled during the differentiation process. In this study, only two DH lines were detected among the 19 regenerated plants (Fig. 2A). The other individuals differed in terms of zygosity and ploidy level from ordinal diploid DH lines (Supplemental Table S2). We could thus readily infer that abnormalities occurred during male gametophyte formation in the interspecific hybrid. We therefore examined anomalies related to male gametophyte formation from two perspectives, genomic zygosity and ploidy level.

### Variations in zygosity

Individuals derived by AC of the interspecific hybrid were divided into three groups on the basis of zygosity: 1) Hom individuals having completely homozygous genomes, 2) Het individuals with complete heterozygosity, and 3) Hom/Het plants having both homozygous and heterozygous genomic regions (Fig. 2A). The first group presumably originated from cases in which the haploid genome of a gamete spontaneously doubled during callus formation or regeneration in an AC-derived rice plant (Rout et al., 2016; Naik et al., 2017). The complete heterozygosity of plants in the second group had two possible causes (Huang et al., 1997): a) callus formation of the F_1_ somatic cells, such as anther wall cells, and b) callus formation occurring in the PMC harboring the paired genomes before the first meiotic division. The third group, which included both homozygous and heterozygous regions, may have emerged after meiotic recombination (Pinson and Rutger, 1993). Tetraploid Hom/Het plants may have been derived from microspores in which the diploid genome was doubled during callus development, while diploid Hom/Het plants may have arisen from microspores formed from callus without genome doubling. AC of rice intraspecific hybrids rarely produced Hom/Het plants, which were derived from diploid microspores (Grewal et al., 2011).

### Meiotic anomalies

In AC of rice, plant differentiation occurs via callus. The most active period of callus formation during pollen development corresponds to the middle to late uninucleate microspore stage. We observed abnormal forms of microspores at the uninuclear stage in the interspecific hybrid, such as microspores that were twice the size of normal ones and fusions of two microspores (Fig. 1C). Flow cytometry and chromosome observations demonstrated that many regenerated plants were tetraploid, triploid, or pentaploid (Fig. 3). These observations suggest that meiotic anomalies of interspecific hybrids lead to insufficient microspore separation and occasional fusion at the tetrad stage. Our genomic analysis revealed that 12 of 19 regenerated plants resulted from abnormalities in division after meiotic recombination (Fig. 4B). Anomalies in meiotic divisions were also observed by immunohistochemical staining for OsCenH3 and α-tubulin (Fig. 4C, D). These meiotic anomalies involved cancellation of either the first or second division, thereby leading to diploid gametophyte generation and subsequent polyploid formation during plant regeneration from callus (Fig. 4) (Jauhar, 2007; De Storme and Geelen, 2013; Han et al., 2018). In some cases, neither the first nor the second division occurred, and the tetraploid gametes were able to develop into callus and differentiate directly into plants. Although a detailed explanation for how triploid and pentaploid plants were generated from AC of the hybrid could not be determined, we were able to deduce the mechanisms associated with the occurrence of tetraploidy based on meiotic anomalies in the interspecific hybrid.

Because most male gametes in F_1_ hybrids between *sat–gla* should decay during pollen development, determination of the genome harbored by each gamete has not been possible. In this study, we demonstrated that genetic characterization of male gametes in hybrid plants between *sat–gla* is feasible by rescuing abortive microspores with AC and allowing them to differentiate into plants. More than a dozen *HS* loci between *sat–gla* can act on male and/or female gametes and, in particular, cause male gametes to become sterile (Koide et al., 2008; Garavito et al., 2010; Kanaoka et al., 2018). Although *HS* genes are widely known to be responsible for HS, our study has clearly shown that meiotic anomalies occur before these genes act (Fig. 5A). Alternatively, meiotic anomalies may also be one of the causes of HS that collapses the gamete genome (Fig. 5A). Future required work includes a detailed analysis of meiotic anomalies occurring in PMCs in hybrids and clarification of the relationship between the mechanism of non-segregation of the first and second divisions and gamete decay.

### Ploidy levels and HS avoidance mechanisms

Among the plants derived from AC, all five plants that produced seeds had tetraploid and heterozygous genomic regions. Four of these five fertile plants were Hom/Het, and one was a completely Het individual. The four Hom/Het plants also had many alleles of *HS* loci as heterozygote. Gametes possessing a killed allele at an *HS* locus will not survive (Sano et al., 1979; Jones et al., 1997). The existence of multiple *HS* loci reduces the number of surviving gametes by one-half per each additional locus. More than a dozen *HS* loci have been found between *sat–gla*, and most of the hybrid gametes are sterile or die (Sano et al., 1979; Sano, 1983, 1990; Doi et al., 1998; Doi et al., 1999; Taguchi et al., 1999; Ren et al., 2006; Zhang et al., 2006; Li et al., 2011; Xu et al., 2014; Yu et al., 2018). In the present study, fertile plants were obtained from a tetraploid with heterozygous *HS* alleles. Except for backcross lines with either parent, we never obtained fertile plants from self-pollinated interspecific F_1_ hybrids between *sat–gla* (Table 1; Supplemental Table S2). Among the 19 regenerated plants from AC, in contrast, we obtained five fertile plants, all of which were tetraploids. Polyploidization may thus be a way to remove the barrier between the two species.

The *HS* genes responsible for the *S1* locus between *sat–gla* have recently been isolated (Xie et al., 2017; Koide et al., 2018; Xie et al., 2019), thus allowing the mechanism of the killer–protector system to be elucidated. In this system, a killer gene is linked to a protector gene that protects gametes from the action of the former (Yang et al., 2012; Ouyang and Zhang, 2013; Zhu et al., 2017; Xie et al., 2019). When a protector gene is present in the same gamete, the killer allele is protected against the killer protein itself. If so, the killer gene at the *HS* locus appears to sporophytically act on other gametes (not encased in the same membrane) that do not have a protector after separation into a tetrad. Tetraploids from *sat–gla* hybrids are likely fertile because three-quarters of diploid gametes from a tetraploid plant contain both killer and protector alleles (Fig. 5B). In contrast, a diploid plant derived from hybrids between *sat–gla* produces haploid gametes, a half of which may contain both killer and protector alleles (Fig. 5B). Although the different killer–protector allele ratios in gametes may reflect the distinct seed fertilities of the tetraploid vs. the diploid, the killer– protector system is not the only explanation for these observations.

### Characteristics of fertile plants obtained from AC

Five lines of fertile tetraploids, #20, #25, #38, #47, and #80, were obtained by AC of the interspecific hybrid of *sat–gla* (Table 1; Supplemental Table S2). The seed sizes of #38 and #80 lines, which produced sufficient seeds, were respectively 1.3 to 1.4 times larger than those of the parental lines and were inherited by the next generation (Supplemental Table S3). Plant heights, flag-leaf lengths, flag-leaf widths, and ligule lengths of plants grown from the seeds of #25, #38, and #80 were superior to the parental traits (Supplemental Table S3). This typical biomass enlargement may have been due to tetraploidization; alternatively, heterosis may have occurred, as the genomes of these strains were heterozygous. In the *sat–gla* diploid F_1_ hybrid, however, the values of these traits were often intermediate between those of the parents, and the tetraploid vigor was thus unlikely the result of heterosis (Supplemental Table S3). Even if heterosis was a factor—given that these tetraploid plants retained heterozygous genomes—the maintenance of heterotic traits in the progeny would be difficult.

In a tetraploid plant with two different alleles at a locus, 10 generations are theoretically required to reduce the proportion of heterozygotes to less than one-quarter of a population; in a diploid plant, this percentage is achieved by the third generation. This characteristic implies that the number of generations during which recombination can take place in a heterozygous tetraploid is much larger compared with a diploid (Pecinka et al., 2011). Tetraploid hybrid plants therefore have the potential to create highly variable allelic combinations by repeated recombination during meiosis.

## Materials and Methods

### Plant materials and AC

The calli derived from AC in this study originated from the same materials obtained by Kanaoka et al.(2018). Interspecific F_1_ hybrid individuals were produced by crossing *O. glaberrima* Steud. with *O. sativa* L. ssp. *japonica*. The seed parent *O. glaberrima* accession IRGC 104038 from Senegal (designated as WK21) was kindly provided by the International Rice Germplasm Center of the International Rice Research Institute (Philippines) and conserved at Kyushu University. Nipponbare (Nip) was used as the pollen parent. Callus induction from AC was carried out according to Kanaoka et al. (2018) and is described as follows. After sterilization with 70% ethanol, panicles with spikelets at the booting stage (uninucleate stage) were incubated at 10°C (low temperature treatment) in the dark for 4 to 10 days. Approximately 70 anthers per dish were plated onto RI-13 callus-induction medium (Woo et al., 1978) prepared in a ø 90 mm × H 15 mm plastic dish. The plated anthers were then cultured at 25°C in the dark for 4 months. Grown calli were transplanted to fresh medium to promote further growth. To induce plant regeneration, calli grown to a diameter of 2 mm were moved to N6 medium (Chu, 1978) and incubated under light conditions at 25°C. When plantlets developed and roots emerged in the medium, the plantlets were transplanted to sterile soil, which included equal amounts of peat moss, vermiculite, and compost. The rice plants were grown under shade conditions in the greenhouse.

### Pollen observation

The anthers for pollen observation were collected based on a distance between the auricles of flag leaf and penultimate leaf. To estimate microspore stages, microspore was collected when then the two auricles were separated by the following distances: −1.0 to +1.0 cm for the uninucleate stage and +2.0 to +6.0 cm for the binucleate state. In addition, mature pollen was collected after heading. These distances were almost all the same among the plant materials used. Collected anthers were fixed with formalin–acetic acid–alcohol fixative and then prepared for microscopic observation. For observation at each microspore developmental stage, anthers were squashed on a microscope slide. After addition of 10 μl acetocarmine or Lugol’s iodine staining solution, the slide was covered with a cover slip and observed to respectively determine the pollen developmental stage or fertility of mature pollen.

### Chromosome counting

For chromosome number estimation, mitosis was observed using cells from root tips of regenerated plant #20, which were pretreated using 2 mM 8-hydroxyquinoline for 2 to 2.5 h at 20°C. After fixation, 1 mm of each root tip was cut off and macerated in enzyme solution consisting of 6.0% (w/v) Cellulase Onozuka RS (Yakult Pharmaceutical, Tokyo, Japan), 6.0% (w/v) Pectolyase Y-23 (Kyowa Chemical Products, Kagawa, Japan), and 75 mM KCl for 60 min at 37°C. The root tips were washed with a drop of distilled water for 5 min on a glass slide. To spread cells, each root tip was thoroughly squashed using a needle with 10 μl ethanol-acetic acid [3:1 (v/v)], and the slide was then flame-dried. The spread cells were stained for 30 min with Giemsa solution (Kanto Chemical Co. INC., Tokyo, Japan) diluted 30 times with Sorensen’s phosphate buffer (pH 6.8). After washing with distilled water, the number of chromosomes was counted under an optical microscope (Olympus BX-50 F, Olympus, Tokyo, Japan).

### Ploidy analysis

Ploidy levels of materials were examined by measuring relative nuclear DNA amounts by flow cytometry as described in Miyashita et al. (2011). Nuclear suspensions obtained by extraction of small pieces of leaf tissue with nuclear extraction buffer (Quantum Stain NA 2A, CytoTechs, Ibaraki, Japan) were filtered through a 30-μm nylon mesh (Partec Celltrics, Lincolnshire, IL, USA). The fluorescent intensity of nuclei stained with DAPI (pH 7.5) was measured using a flow cytometer (Partec PA, Partec GmbH, Münster, Germany). The ploidy level of each examined individual was estimated using the fluorescent intensity of diploid tissue as a standard.

### Genotyping

PCR detection of polymorphisms between WK21 and Nip was based on comparison of their complete genome sequences. The complete genome sequence of Nip was obtained from IRGSP-1.0 (RAP-DB), while that of WK21 was sequenced and deposited into the DDBJ under accession number DRS049718. Genomic DNA of regenerated plants from WK21/Nip F_1_ individuals were extracted from mature leaves of well-grown regenerants. For genotyping of regenerated plants, we used 57 markers designed using SSR or InDel polymorphisms between WK21 and Nip (Supplemental Fig. 2). Among the 57 markers, 22 were randomly distributed on each of 12 chromosomes (McCouch et al., 2002), and 24 were located near the centromere of each chromosome. Each centromere location was based on the Rice Genome Annotation Project database (http://rice.plantbiology.msu.edu/cgi-bin/gbrowse/rice/). In addition, 12 markers linked to *HS* loci were used to test zygosity. PCR amplifications for genotyping were performed using GoTaq Green Master Mix (Promega, Madison, WI, USA), with the resulting products subjected to 3% agarose gel electrophoresis (Supplemental Fig. S3). Three genotyping analyses were independently performed.

### Immunohistochemical staining

Samples were soaked for 20 min in a fixative consisting of microtubule-stabilizing buffer (5 mM PIPES, 0.5 mM MgSO_4_, and 0.5 mM EGTA, pH 7.0) containing 3% (w/v) paraformaldehyde and 0.1% (v/v) Triton X-100 and then rinsed twice in 1× PBS buffer for 10 min. In the primary reaction, two primary antibodies were used: anti-OsCenH3 rabbit antibody and anti-α-tubulin mouse antibody (T6199, Sigma-Aldrich, St. Louis, MO. USA) (Nagaki et al., 2004). A primary antibody solution containing the two antibodies was diluted 200 times with a blocking buffer [0.4 M Tris-HCl (pH 7.5), 3.5% (w/v) NaCl, and 2% (w/v) BSA]. Fixed anthers were gently dissected on a glass slide using tweezers. Cells from the dissected anthers were suspended in 20 μl of 1× PBS, and covered with a coverslip, and then stored in a freezer (−80°C). After freezing, the coverslip was removed, 100 μl of the primary antibody solution was applied, and the solution was covered by a piece of parafilm (55 × 26 mm) to spread the solution. The samples were placed in a moisture chamber to prevent drying and kept at 4°C for 14 h. After the primary reaction, the samples were rinsed three times with 1× PBS for 10 min. Two secondary antibodies were used: Alexa Fluor 488-labeled anti-mouse antibody (#A-11001: Invitrogen, Carlsbad, CA, USA) and Alexa Fluor 555-labeled anti-rabbit antibody (#A20739: Invitrogen). The secondary antibody solution was diluted 200 times with the same blocking buffer used in the primary reaction. After the washing, the PBS buffer was removed from the slides, and then 100 μl of secondary antibody solution was applied, and the solution was covered by a piece of parafilm. The slides were placed in a moisture chamber and incubated at 37°C for 1 h. After the secondary reaction, the samples were rinsed using the same procedure applied after the primary reaction and then dried. To stain DNAs with minimal fading, 20 μl of ProLong Diamond Antifade Mountant with DAPI (Invitrogen) was applied to each slide before observation.

## Acknowledgments

We are grateful for the help of Dr. S. Ishiguro, Mr. Y. Ota, and Ms. R. Iwashiro (Laboratory of Plant Breeding, Hokkaido University) in conducting this study. We also thank Dr. N. Ohmido (Graduate School of Human Development and Environment, Kobe University) for valuable technical advice. We thank Edanz Group (www.edanzediting.com/ac) for editing the English text of a draft of this manuscript.

